# Enhanced flux prediction by integrating relative expression and relative metabolite abundance into thermodynamically consistent metabolic models

**DOI:** 10.1101/481499

**Authors:** Vikash Pandey, Noushin Hadadi, Vassily Hatzimanikatis

## Abstract

The ever-increasing availability of transcriptomic and metabolomic data can be used to deeply analyze and make ever-expanding predictions about biological processes, as changes in the reaction fluxes through genome-wide pathways can now be tracked. Currently, constraint-based metabolic modeling approaches, such as flux balance analysis (FBA), can quantify metabolic fluxes and make steady-state flux predictions on a genome-wide scale using optimization principles. However, relating the differential gene expression or differential metabolite abundances in different physiological states to the differential flux profiles remains a challenge. Here we present a novel method, named REMI (<u>R</u>elative <u>E</u>xpression and <u>M</u>etabolomic <u>I</u>ntegrations), that employs genome-scale metabolic models (GEMs) to translate differential gene expression and metabolite abundance data obtained through genetic or environmental perturbations into differential fluxes to analyze the altered physiology for any given pair of conditions. REMI is the first method that integrates thermodynamics together with relative gene-expression and metabolomic data as constraints for FBA. We applied REMI to integrate into the *Escherichia coli* GEM publicly available sets of expression and metabolomic data obtained from two independent studies and under wide-ranging conditions. The differential flux distributions obtained from REMI corresponding to the various perturbations better agreed with the measured fluxomic data, and thus better reflected the different physiological states, than a traditional model. Compared to the similar alternative method that provides one solution from the solution space, REMI was also able to enumerate several alternative flux profiles using a mixed-integer linear programming approach. Using this important advantage, we performed a high-frequency analysis of common genes and their associated reactions in the obtained alternative solutions and identified the most commonly regulated genes across any two given conditions. We illustrate that this new implementation provides more robust and biologically relevant results for a better understanding of the system physiology.

**Author Summary:** The recent advances in omics technologies have provided us with an unprecedented abundance of data spanning genomes, global gene expression, and metabolomes. Though these advancements in high-throughput data collection offer an excellent opportunity for a more thorough understanding of metabolic capacities of a wide range of species, they have caused a considerable gap between “data generation” and “data integration.” reconstructed model to predict the observed physiology, e.g., growth phase through omics data integration. In this study, we present a new method named REMI (<u>R</u>elative <u>E</u>xpression and <u>M</u>etabolomic <u>I</u>ntegrations) that enables the co-integration of gene expression, metabolomics and thermodynamics data as constraints in genome-scale models. This not only allows the better understanding of how different phenotypes originate from a given genotype but also aid to understanding the interactions between different types of omics data.

## Introduction

The turnover rates of metabolites through a pathway are called fluxes, and genome-wide intracellular metabolic fluxes are the ultimate regulator of cellular physiology. Perturbations on this normal physiology, such as those that occur in a disease state, directly influence the metabolic fluxes. The well-established experimental approach for determining these metabolic fluxes is ^13^C metabolic flux analysis, though this experimental technique that directly measures metabolite levels is costly and time-consuming, such that computational tools for flux prediction have become a very popular alternative. Genome-scale metabolic models (GEMs), which essentially associate an organism’s genotype with its phenotype, integrate genomic information with known information about metabolite levels to comprehensively describe an organism’s metabolism [1]. These models can predict metabolic fluxes, growth rates, or the fitness of gene knockouts using constraint-based approaches, which mainly require the knowledge of network stoichiometry that is available from the annotated genome sequences and metabolic pathway databases. One of the most routinely used constraint-based approaches is flux balance analysis (FBA), which relies on the stoichiometry and optimization principles to predict the steady-state metabolic flux distribution according to an objective function in a given metabolic network [2]. Due to network complexity, FBA commonly results in a span of alternative optimal solutions indicating different flux distributions with the same objective value rather than a unique steady-state flux distribution profile, and then selects one of these solutions at random to present back to the user, which is a major limitation of this method. To remedy this, it has been shown that integrating additional layers of constraints, such as thermodynamics, can effectively reduce the overall solution space of feasible flux distributions in an organism to limit the number of alternative solutions [3, 4].

With the growing availability of high-throughput data for different organisms under a wide range of genetic or environmental perturbations, GEMs became popular because of their ability to incorporate omics data as additional regulatory constraints for FBA problems. Because GEMs associate a genotype with a phenotype, it is essential to understand that a single genome can result in thousands of different physiologies through different regulatory mechanisms. Therefore, the integration of static snapshots of the metabolism, obtained from transcriptomic and metabolomic data, provides more biologically relevant constraints for the system and helps to increase the precision of the flux prediction, therefore better deducing the observed physiology. However, despite the high number of methods that have been introduced in recent years for the integration of omics data into constraint-based metabolic models, the enhanced prediction of flux profiles using omics data, particularly in cases using multi-omics data, is still far from being resolved. Recently, these methods, their scopes, and limitations were extensively reviewed [5], and the authors concluded that using gene-expression data does enhance flux predictions, though they inferred that the accurate predictions of the physiology is not achievable with the available reviewed methods.

The existing methods for integrating gene-expression data into GEMs can be classified into two categories with the first relying on the integration of absolute gene-expression data into GEMs. This includes techniques such as gene inactivity moderated by metabolism and expression (GIMME; [6]) and the use of continuous and discrete formulations to find a flux distribution that is consistent with given context-specific gene-expression data, including integrative metabolic analysis tools (iMAT; [7, 8]) [5, 9-11]. However, the assumption that absolute gene-expression data can be directly correlated with flux values is questionable and might not hold true for all genes. Moreover, these methods require user-defined thresholds to identify and categorize the expression levels of metabolic genes (high, moderate, or low expression), and the results are sensitive to the set thresholds. These drawbacks motivated the development of (ii) the second class of methods, which integrate the relative gene-expression data while aiming to maximize the correlation between differential changes in gene-expression and reaction fluxes. The underlying assumption for this class of methods is that the relative changes in gene expression between two conditions correlate with the resulting differential flux profiles [12, 13].

The increasing availability and quality of metabolomic data have promoted the development of methods that can be integrated into GEMs to refine model reconstruction, to reduce the solution space of feasible fluxes, and to better predict the physiological state of a system. These methods, their scope, and their limitations have been reviewed by Töpfer *et al.* [14]. One of these methods, thermodynamic-based flux balance analysis (TFA), integrates the absolute metabolite concentration data into GEMs, as the metabolite concentrations are intrinsically associated with the Gibbs free energy of metabolic reactions [3, 4]. Another available method is gene inactivation moderated by metabolism, metabolomics, and expression (GIM3E), an extension of the GIMME algorithm with added metabolomic data in addition to gene-expression data [15]. However, this method only considers the presence/absence of metabolites to refine the model, therefore preventing a full utilization of the quantitative metabolomic data. A time-resolved expression and metabolite-based prediction of flux values, named TERM-FLUX, integrates time-series expression and metabolomic data, and predicts flux distribution for a given time point t. [16]. However, the application of TERM-FLUX is limited to studies with time-series data, which are not widely available. More recently, a method for the integration of relative metabolite levels for flux prediction, iReMet-flux, has been introduced to predict differential fluxes at the genome-scale [17], and it requires an assessment of the differential changes of all existing metabolites in a GEM. This limits its application, as metabolomic data are mostly measured not at a genome-wide level but rather for only a few metabolites in a system.

For multi-omic data, methods have recently been introduced for integrating different layers of data, such as genomic, transcriptomic, proteomic, and fluxomic, into metabolic models [18] or multi-scale models [19]. However, a method that couples the thermodynamic constraints into GEMs with relative transcriptomic and metabolomic data is not yet available.

To address this deficiency, we herein propose a novel method, termed **R**elative **E**xpression and **M**etabolite **I**ntegration (REMI), to integrate relative expression and relative metabolite abundance data into thermodynamically curated GEMs. REMI allows for gene-expression, metabolite abundance, and thermodynamic data to be integrated into a single framework, then uses optimization principles to maximize the consistency between the differential gene-expression levels and metabolite abundance data and the estimated differential fluxes and thermodynamic constraints. We demonstrate that REMI’s ability to integrate different layers of constrictive data significantly reduces the solution space of feasible fluxes. REMI also extensively enumerates alternative optimal and sub-optimal solutions, bringing a robustness and flexibility to the flux distribution analysis. We applied REMI to an *E.coli* GEM to estimate the central carbon metabolism intracellular flux measurements that were determined by ^13^C metabolic flux analysis (^13^C-MFA) and were provided by two independent experimental studies [20, 21]. Using transcriptomic and metabolomic data from the different experimental conditions, we observed a remarkable correlation between experimental fluxes and the predicted fluxes. Comparing REMI’s predictions with a similar method (GX-FBA [12]), we also show that REMI has on average a 32% higher Pearson correlation coefficient (r = 0.79) indicating a more precise exploration of organismal metabolism under wide-ranging conditions.

## Results and Discussion

We designed REMI as the first method to integrate relative gene-expression and metabolite abundance data into thermodynamically curated GEMs, significantly reducing the solution space of feasible fluxes to provide results that are better at predicting cell physiologies closer to the experimental observations than can be reached using existing methods. The REMI framework was applied to integrate the *E. coli* transcriptomic and metabolomic data obtained from two studies under 8 [20] and 3 [21] different conditions into the thermodynamically curated *E. coli* GEM iJO1366 and to estimate the differential steady-state fluxes. We call the data and information from [20] “Dataset A” and data and information from [21] “Dataset B”. We formulated different optimization models which hierarchically integrated different combinations of available data to investigate the effectiveness of multi-omic data integration in reducing the metabolic flexibility of the provided solutions. REMI-TGex is an integrated model obtained by incorporating relative gene-expression data into a thermodynamically constrained model, which is represented by iJO1366 in this work. Furthermore, we integrated relative metabolite concentration data into the REMI-TGex model to produce REMI-TGexM and compared experimentally measured fluxes with the steady-state flux prediction results of REMI-TGex and REMI-TGexM. We also compared our prediction results with those of the previously existing GX-FBA [12], though as this method does not employ thermodynamic constraints, we used REMI to incorporate gene-expression data lacking thermodynamics constraints into *E. coli* GEM iJO1366 (REMI-Gex). The comparison of REMI-Gex and REMI-TGex highlights the significance of the thermodynamic constraints in reducing the solution space of flux analysis. We also performed some studies with only metabolite changes with thermodynamic constraints (REMI-TM) and without thermodynamic constraints (REMI-M).

### Consistency score and enumeration of alternative solutions

The underlying assumption of the REMI method is that the perturbation of gene-expression and metabolite levels influences the flux levels in the metabolic network. To this end, REMI maximizes the consistency between relative experimentally observed changes in gene expression and metabolite changes with the flux levels (the objective function of the REMI constraint-based method). The maximum consistency is then calculated as an integer number, called the maximum consistency score (MCS). This represents the maximum number of constraints that can be incorporated into a FBA model from a given set of constraints (gene-expression or metabolite abundance levels) while ensuring that the model still achieves the required metabolic functionalities and remains feasible. MCS is a unique number, however, in that the complex nature and interconnectivity of metabolic networks can result in several alternative solutions for a given MCS, meaning that numerous combination of different constraints from the input data could result in the same MCS. The theoretical maximum consistency score (TMCS) indicates the number of genes (or metabolites or both) with available experimental data that can potentially be integrated into the model, and MCS indicates the number of these available constraints that could be consistently integrated into the model.

### Case study I: REMI analysis of the Dataset A with gene-expression and metabolomic data

We first applied REMI to the integration of eight datasets from Ishii *et al.* [20], which included genome-wide transcriptomics together with some metabolomic data obtained for one reference condition and seven different conditions or mutations, into an *E. coli* model. After integrating the gene-expression data of each condition into the model and comparing it with the reference model, we computed TMCSs, MCSs, and the number of alternative solutions for the REMI-Gex method (without thermodynamic constraints) and the REMI-TGex method (with thermodynamic constraints). In contrast to other methods, REMI finds all possible alternative solutions of a given maximum consistency score, which involves all possible combinations of the given set of constraints that always result in a feasible model. These alternative solutions provide flexibility in the biological interpretation of the results as they are equally consistent with the provided experimental data (applied as constraints to the model). Note that in the GEM analysis, the alternate flux solutions are conventionally considered as equivalent phenotypic states [22]. In this study, however, alternative solutions represent the equivalent states of the maximum consistency between gene-expression (or metabolite abundance or both) data and the flux levels. Therefore, each feasible alternative solution provides an opportunity to analyze and interpret the given phenotypic state based on the condition-specific omics data, from a different standpoint.

We further integrated the available metabolomics measurements into the *E. coli* model using REMI-TGexM and obtained the MCS for the integrated metabolites as well as the global maximum consistency score (GMCS), which encompasses both genes and metabolites (Table 1). Although different pairs of conditions showed a very close TMCS variability across the seven case studies based on gene-expression data integration (mean = 103.6, standard deviation [sd] = 0.5) and on metabolic data (mean = 4.7, sd = 0.5), the MCS significantly varied across the four REMI methods: REMI-Gex (mean = 58.7, sd = 4.1), REMI-TGex (mean = 49.7, sd = 3.5), REMI-GexM (mean = 63.3, sd = 4), and REMI-TGexM (mean = 54.3, sd = 3.3) (Table 1). As Table 1 shows, the gene-expression and metabolite abundance constraints for the deregulated metabolites were consistent across the conditions. Therefore, the REMI-TM and REMI-GexM consistency scores sum up to the REMI-TGexM consistency score. This means that there is no conflict between the gene-expression and metabolite abundance data and that they can be co-integrated without confronting each other. However, the number of enumerated alternative solutions highly differs across the conditions in all four methods: REMI-Gex (mean = 80.6, sd = 80.3), REMI-TGex (mean = 104.6, sd = 168.5), REMI-GexM (mean = 156.1, sd = 241.8), and REMI-TGexM (mean = 25.1, sd = 26.4) (Table 1), which suggests that the numbers of alternative solutions are condition-specific, as expected. As shown in the Table 1, wherever the sd is very high, for example sd=241.8 in REMI-GexM for the rpe *vs* Ref case, we observe a high number of alternative solutions (n=735 in this case). Different conditions (mutations) alter the cell metabolism differently, leading to different levels of metabolic adaptations and metabolic flux rerouting. Hence, we speculate that the differences in flux rerouting across conditions results in differences in the numbers of alternative solutions across the seven relative conditions. Note that for REMI-TM and consequently for REMI-M, the constraints for all the deregulated metabolites were consistently integrated into the model, so we found only one solution for the REMI-TM models without any alternative solution.

**Table 1.** Maximum consistency score and the number of alternative solutions for different models. The reference refers to the wildtype growth rate of 0.2/hour. TMCS, theoretical maximum consistency score; MCS, maximum consistency score; # Alt, number of alternatives; HFC, high frequency constraint. Metabolites represent differentially regulated metabolite between two conditions. SD represents the standard deviation across comparisons.

| Comparisons | Genes | REMI-Gex |  |  | REMI-TGex |  |  | Metabolites |  | REMI-GexM |  |  | REMI-TGexM |  |  |
| --- | --- | --- | --- | --- | --- | --- | --- | --- | --- | --- | --- | --- | --- | --- | --- |
|  | TMCS | MCS | # Alt | HFC | MCS | # Alt | HFC | TMCS | MCS | GMCS | # Alt | HFC | GMCS | # Alt | HFC |
| pgm vs Ref | 104 | 56 | 11 | 50 | 49 | 8 | 45 | 5 | 5 | 61 | 12 | 55 | 54 | 8 | 50 |
| gapC vs Ref | 104 | 60 | 16 | 56 | 48 | 16 | 44 | 5 | 5 | 65 | 16 | 61 | 53 | 16 | 49 |
| zwf vs Ref | 104 | 62 | 8 | 59 | 54 | 4 | 52 | 5 | 5 | 67 | 8 | 64 | 59 | 4 | 57 |
| rpe vs Ref | 104 | 58 | 67 | 48 | 49 | 512 | 40 | 5 | 5 | 62 | 735 | 52 | 53 | 4 | 46 |
| pgi vs Ref | 103 | 59 | 236 | 50 | 49 | 64 | 43 | 4 | 4 | 63 | 96 | 56 | 53 | 16 | 49 |
| wt5 vs Ref | 103 | 65 | 160 | 58 | 55 | 80 | 49 | 4 | 4 | 69 | 160 | 62 | 59 | 80 | 53 |
| wt7 vs Ref | 103 | 51 | 66 | 43 | 44 | 48 | 38 | 5 | 5 | 56 | 66 | 48 | 49 | 48 | 43 |
| Mean | 103.6 | 58.7 | 80.6 | 52.0 | 49.7 | 104.6 | 44.4 | 4.7 | 4.7 | 63.3 | 156.1 | 56.9 | 54.3 | 25.1 | 49.6 |
| SD | 0.5 | 4.1 | 80.3 | 5.4 | 3.5 | 168.5 | 4.5 | 0.5 | 0.5 | 4.0 | 241.8 | 5.4 | 3.3 | 26.4 | 4.2 |

### Alternative solutions and consistency scores of REMI-TGex and REMI-TGexM

For the metabolomic integration, the GMCS was higher in the REMI-TGexM models compared to REMI-TGex because in REMI-TGexM, the GMCS was computed based on both relative metabolite (Table 1; Metabolites) and relative gene-expression levels (Table 1; Genes), whereas the MCS for the REMI-TGex model was computed based on only relative expression levels. We further investigated the consistency between gene-expression and metabolomic data and whether the data contradicted each other in certain scenarios. All the available experimental metabolomic data (Table 1; TMCS and MCS) were integrated using the REMI-TGexM method for the pgm *vs* Ref, gapC *vs* Ref, zwf *vs* Ref, wt5 *vs* Ref, and wt7 *vs* Ref comparisons. We observed that the number of alternative solutions for these five cases was identical between REMI-TGexM and REMI-TGex. This implies that the relative expression constraints and the relative metabolite constraints were not contradictory for these five cases. However, in rpe *vs* Ref and pgi *vs* Ref, all the metabolic data were integrated in the model, but the number of alternative solutions differed (and in the case of rpe *vs* Ref was noticeably reduced) between REMI-TGexM and REMI-TGex. To see if this indicated a contradiction, further investigation into the alternative solutions revealed that in the rpe *vs* Ref and wt5 *vs* Ref comparisons, REMI-TGexM and REMI-TGex have the same set of constraints, which means that the constraints from metabolomics and expression data were not contradictory. However, we found that the metabolomics integration resulted in a reduction in the number of alternative solutions (Table 1). We hypothesized that further integration of metabolomics (on the top of the gene-expression constraints) imposed a flux rerouting in the metabolic network.

### High-frequency constraint (HFC) analysis

As REMI allows enumerating all the possible alternative solutions for a given consistency score, we further interrogated the alternative solutions by High-frequency constraint (HFC) analysis.

The results of this analysis indicate the core constraints that consistently operate in all the alternative solutions (the constitutive part of all solutions). Meaning that such core constraints certainly perturb fluxes within each pair of conditions. Therefore, these constraints could potentially be the indicators of the regulators of the condition-specific metabolism, which assist biologist in determining which metabolic subsystems to deregulate or to mutate. We believe that the capability to analyze and identify these regulators is a key advantage of REMI.

As shown in the Table 1, the computed HFCs differ across conditions for all four cases: REMI-Gex (mean = 52, sd = 5.4), REMI-TGex (mean = 44.4, sd = 4.5), REMI-GexM (mean = 56.9, sd = 5.4), and REMI-TGexM (mean = 49.6, sd = 4.2). Constraints that were common amongst all the alternative solutions, indicating key regulators, were the potential candidates for further investigations. After analyzing HFCs across conditions and between the four cases, we found that a reaction catalyzed by glycolate oxidase (GLYCTO4) from the alternate carbon metabolism and another reaction from the murine recycling pathway (MDDEP4pp) were always deregulated in the pgm, gapC, zwf, rpe, pgi, and wt7 conditions. These reactions are likely key regulators of mutation in *E. coli* because they were found to be deregulated in all mutant conditions.

### REMI-Gex vs REMI-TGex: flux variability analysis to investigate the influence of thermodynamic constraints

To study the effect of thermodynamics on the model, we compared the reduction in solution space for the predicted flux profiles from the REMI-TGex and REMI-Gex methods when coupled with the gene-expression data (Table 1). The MCS was consistently reduced in the REMI-TGex model compared to REMI-Gex for all pairs of conditions, as REMI-TGex eliminates flux solutions that are not thermodynamically feasible.

To better illustrate the positive influence of thermodynamic constraints in reducing the solution space, we show the example of pgm *vs* Ref as a case study, where we obtained MCS = 56 in REMI-Gex and MCS = 49 in REMI-TGex (Table 1). First, we enforced the models to satisfy any given consistency score (56 and 49 in this example) by adding a new constraint, which would further allow us to perform conditional FVA. Then, we performed the FVA that satisfies the consistency score (MCS = 56) in REMI-Gex and the consistency score (MCS = 49) in REMI-TGex. Comparing the FVA results of REMI-Gex and REMI-TGex revealed that there exist 45 reactions in REMI-Gex that operate in a thermodynamically infeasible direction and which also contribute to the MCS = 56. The flux ranges of these reactions are shown in Table S1 and indicate that the TGex method is indeed eliminating the infeasible solutions to enrich for more relevant results. For more clarification, two reactions out of the 45 are shown as examples in Figure 1. As expected, the flux ranges for these reactions are less flexible for the REMI-Gex (MCS = 56) compared to the REMI-TGex (MCS = 49), which confirms some extent of the thermodynamic infeasibility in the REMI-Gex predictions as infeasible flux ranges directly indicate the model infeasibility. On the other words, if we integrate thermodynamic constraints to the model and allow the consistency score (MCS=56) then the model certainly generates infeasible solutions. To investigate whether the higher consistency score caused thermodynamic infeasibility in the REMI-Gex, we performed a FVA of REMI-Gex while forcing lower consistency scores (MCS = 49 and 10). We found that the flux ranges of reactions became more flexible at lower consistency scores in the REMI-Gex model compared to the REMI-TGex model (Figure 1), indicating that if both REMI-TGex and REMI-Gex have the same consistency scores, the REMI-Gex cannot allow thermodynamic infeasibility. In contrast, if the consistency score is higher in the REMI-Gex compared to the REMI-TGex, then it leads to thermodynamic infeasibility. The same results were obtained for all other reactions (Table S1).

**Figure 1:**
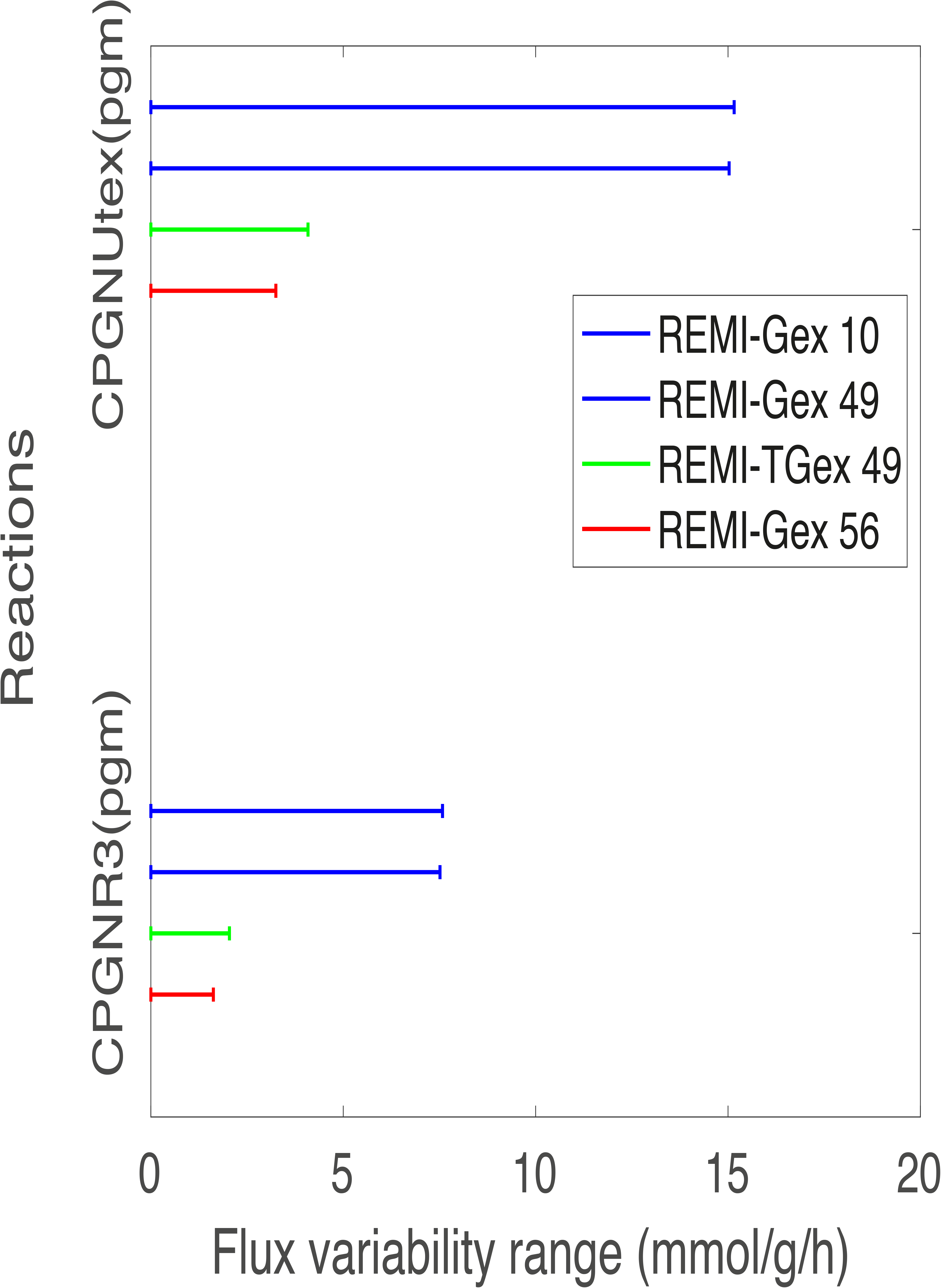
Thermodynamic infeasibility test for two example reactions, CPGNR3 [pgm] and CPGNUtex [pgm]. The bars represent the flux variability range, with a longer bar indicating greater flexibility of feasible flux ranges. Flux ranges are calculated based on the flux variability analysis of the REMI-TGex and REMI-Gex models at different consistency scores. For more clarification, two reactions out of the 45 are shown as examples, where it can easily be seen that there is a much greater variability in the Gex-10 and Gex-49 models by the length of the bar. This indicates that these two models are more flexible compared to the TGex-49 model. This means an equal consistency score (Gex-49) or a lower consistency score (Gex-10) in the absence of thermodynamic constraints (instead of Gex-56) provides a greater flexibility compared to TGex-49.

### Case study II: REMI analysis of the Dataset B with gene-expression and fluxomic data

To further benchmark REMI with the available experimental data, we used a second data set (2 overexpression compared to the ref condition) from an independent study where the role of metabolic cofactors, such as NADH and ATP in different aspect of metabolism is studied by overexpressing NADH oxidase and the soluble F1-ATPase in *E. coli* [21]. REMI integrated the gene-expression data from Holm *et al.* [21] into the *E. coli* model, and a summary of the results is shown in Table 2. Like the previous analysis, we observed a reduction in the MCS value within REMI-TGex as compared to REMI-Gex, as REMI-Gex satisfies fluxes that were not thermodynamically feasible. The number of alternative solutions highly differs between NOX overexpression and ATPase overexpression for both REMI-TGex and REMI-Gex, which is likely due to the condition-specific regulations (NOX *vs* ATPase overexpression) that do not necessarily involve the same set of deregulated genes.

**Table 2.** Maximum consistency score and alternatives for REMI-Gex and REMI-TGex in second *E. coli* study. “Ref” represents the wildtype, and ATPase overexpression and NOX overexpression are relative to the wildtype. TMCS, theoretical maximum consistency score; MCS, maximum consistency score; # Alt, number of alternatives; HFC, high frequency constraint.

| Comparisons | TMCS | REMI-Gex |  |  | REMI-TGex |  |  |
| --- | --- | --- | --- | --- | --- | --- | --- |
|  |  | MCS | # Alt | HFC | MCS | # Alt | HFC |
| NOX vs Ref | 202 | 64 | 48 | 58 | 61 | 16 | 57 |
| ATPase vs Ref | 200 | 75 | 384 | 66 | 65 | 96 | 58 |

### Bidirectional reaction analysis with and without thermodynamics

To investigate the influence of thermodynamic constraints on flux ranges, we identified the overlapping constraints (HFCs) across all the alternative solutions and then enforced them to be active to build the most consistent model. An active HFC satisfies differential gene expression (or metabolite levels) between two conditions form a given experimental data. Thus, for each condition, we build the most consistent model despite having many alternatives. We next performed FVA on the REMI-Gex and REMI-TGex models. As REMI is based on pair-wise relative constraints (for two conditions) and builds two models that are then compared as opposed to modifying one solution based on a given condition, we obtained two FVA solutions, i.e. one for each condition. We identified less bidirectional reactions (BDRs) in the REMI-TGex case compared to the REMI-Gex case (Table 3), which means that thermodynamic constraints reduce the solution space and consequently the number of BDRs. This is consistent with the fact that thermodynamic constraints eliminate infeasible reaction directionalities. The number of BDR reductions differs across conditions, and we identified the highest BDR reduction for the rpe *vs.* Ref case and the lowest BDR reduction for the NOX vs. Ref case, which therefore indicates more reduction in the feasible flux solution space in the rpe *vs.* Ref case compared to the NOX vs. Ref case. For the all comparisons, we found a further reduction in BDRs upon the integration of relative metabolomic data into the REMI-TGex model. In most of the cases, we found a similar decrease in BDRs, which means that the metabolomic data further constrained the solution space. Except for the wt7 *vs* Ref case, we observed a decrease in BDRs for all cases that were constrained by metabolites and expression data together (GexM) as compared to only expression (Gex) data. Unexpectedly and unlike all the other cases, by incorporating metabolomics data for the wt7 *vs* Ref case, we found an increase of one reaction in the BDRs. This suggests that for the wt7 *vs* Ref case the integration of gene expression and metabolites reroutes fluxes through the metabolic networks differently compared to other cases. As expected, we consistently find a reduction in BDRs for the REMI-TM model (thermodynamics and relative metabolomics) in compared to without thermodynamics (the REMI-M model). This is in agreement with the fact that integrating thermodynamic constraints into a model eliminates infeasible reaction directionalities and consequently the flux feasible solution space.

**Table 3.** Uni- and bidirectional reactions for the REMI-Gex and REMI-TGex models in the second *E. coli* study. Table entries are in the form of S (n1,n2), where as S represents the sum of n1 and n2, n1 is the number of bidirectional reactions for the mutant model, and n2 represents bidirectional reactions for the reference model.

| Comparisons | BDRs in REMI-Gex (model1, model2) | BDRs in REMI-TGex (model1, model2) | BDRs in REMI-TGexM (model1, model2) | BDRs reduction (GeX-TGex) | BDRs in REMI-GexM (model1, model2) | BDRs in REMI-TM (model1, model2) | BDRs in REMI-M (model1, model2) |
| --- | --- | --- | --- | --- | --- | --- | --- |
| pgm vs Ref | 222 (109,113) | 188 (92,96) | 186 (93,93) | 34 | 209 (103,106) | 204 (102,102) | 224 (112,112) |
| pgi vs Ref | 247 (132,115) | 205 (103,102) | 169 (78,91) | 42 | 209 (103,106) | 204 (102,102) | 224 (112,112) |
| gapC vs Ref | 201 (103,98) | 158 (77,81) | 156 (76,80) | 43 | 195 (100,95) | 205 (103,102) | 224 (112,112) |
| zwf vs Ref | 226 (114,112) | 200 (102,98) | 192 (98,94) | 26 | 220 (110,110) | 204 (102,102) | 224 (112,112) |
| rpe vs Ref | 260 (135,125) | 201 (99,102) | 169 (89,80) | 59 | 224 (112,112) | 198 (100,98) | 218 (110,108) |
| wt5 vs Ref | 223 (112,111) | 202 (102,100) | 200 (100,100) | 21 | 223 (111,112) | 204 (102,102) | 224 (112,112) |
| wt7 vs Ref | 223 (112,111) | 198 (101,97) | 194 (100,94) | 25 | 224 (112,112) | 204 (102,102) | 224 (112,112) |
| NOX vs Ref | 208 (104,104) | 188 (96,92) |  | 20 |  |  |  |
| ATPase vs Ref | 229 (116,113) | 202 (101,101) |  | 27 |  |  |  |

### Relative flexibility analysis with and without thermodynamics

To further illustrate the positive influence of thermodynamic constraints in reducing the feasible solution space, we performed a relative flexibility (Materials and Methods) analysis using the REMI-TGex and REMI-Gex methods. To perform a relative flexibility analysis, a reference model is compared to a target model to investigate the relative flux reduction. For a reference, we used the iJ01366 model without integrating any data, meaning that the reference model implies only mass balance constraints. We took the pgi *vs.* Ref case as an example to demonstrate the average relative flexibility (ARF) reduction at a global (e.g. all reactions) level as well as at the subsystem level.

For the pgi *vs.* Ref case, we found a 10%, 20%, 50%, 77%, and 80% reduction in the global ARF in RMI-M, REMI-Gex, REMI-TGex, REMI-GexM, and REMI-TGexM models compared to the reference model, respectively (Figure 2a). We found 40% and 80% more reduction in the global ARF for the REMI-TGex and REMI-TGexM models compared to REMI-Gex (Figure 2a), which was expected as the REMI-TGex and REMI-TGexM models are more constrained by thermodynamic and metabolomic data compared to REMI-Gex. We further analyzed the ARF at the subsystem/pathway level to investigate the reduction in ARF for each specific subsystem using the REMI-TGex and REMI-Gex methods. Consistently, each subsystem for the REMI-TGexM and REMI-TGex models was more reduced than REMI-Gex (Figure 2b). For REMI-TGex and REMI-TGexM, we observed a remarkable ARF reduction in the glycerophospholipid metabolism, lipopolysaccharide biosynthesis, murein recycling and biosynthesis, and the biomass and maintenance function subsystems. We further performed the same analysis for the pgi *vs.* Ref, rpe vs. Ref, pgm *vs.* Ref, wt5 *vs.* Ref, wt7 *vs.* Ref, NOX *vs.* Ref, and ATPase *vs.* Ref data (Figure S1). We found a similar reduction in ARF for REMI-TGex and REMI-TGexM compared to REMI-Gex for the cases of gapC *vs.* Ref and zwf *vs.* Ref and found a small reduction in pgi *vs.* Ref and rpe *vs.* Ref (Figure S1). We identified a remarkable reduction in ARF (more than 90%) across all the comparisons using the REMI-TGexM method for the glycerophospholipid metabolism, murein recycling, and lipopolysaccharide biosynthesis/recycling subsystems (Table S2). This suggests that these subsystems are more perturbed based on our available gene-expression and metabolite level data, which indicates that they might be key regulator pathways for the studied mutations.

**Figure 2.**
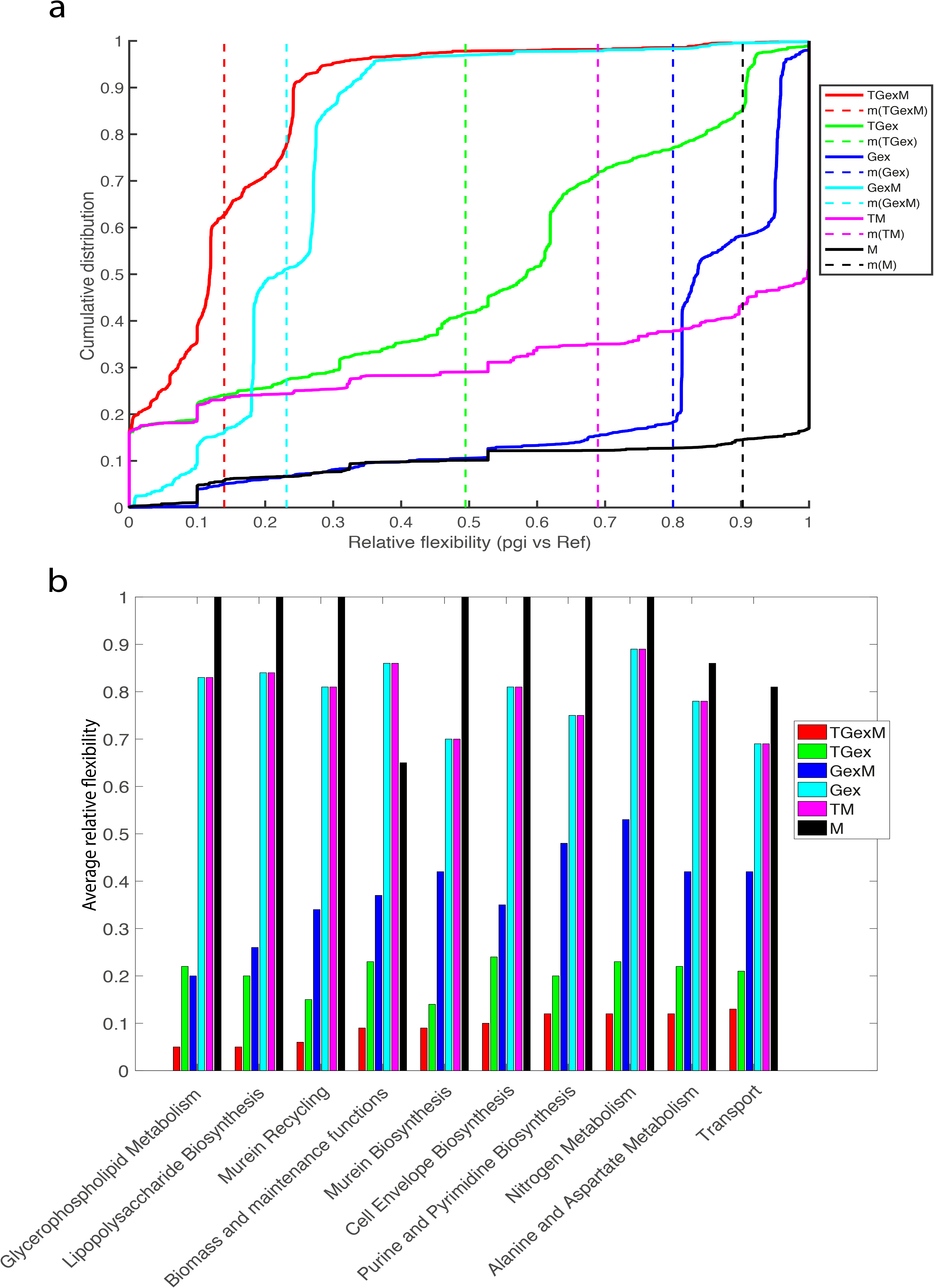
Relative flexibility for the pgm mutant vs. reference case. a) Cumulative relative flexibility of the reactions. Solid lines represent the distribution of relative flexibility of reactions and dotted lines represent the average relative flexibility of reactions. Average relative flexibility for models with thermodynamic (TGexM, TGex, and TM) are smaller compared to their respective in models without thermodynamic (GexM, Gex, and M). This designates the elimination of flux solution space due to thermodynamically infeasible reaction directionality. Interestingly, we found the most reduction in TGexM model which is the most constrained model by three data types. b) The average relative flexibility is shown for the top ten (according to TGexM) metabolic subsystems. Similar to part a, thermodynamically constrained models show a bigger reduction in the feasible flux solution space at the metabolic subsystem level.

### Comparing predictions of different REMI models with the GX-FBA method

To demonstrate the efficacy of the REMI methods in reducing the solution space and therefore predicting flux profiles close to the experimental measurements, we compared the flux predictions of the REMI-Gex, REMI-TGex, and REMI-TGexM methods with those of the alternative, previously used GX-FBA method and compared both methods to the available experimental measured fluxes from ^13^C experiments. To implement the GX-FBA method, we integrated the relative gene-expression datasets into the iJO1366 model using GX-FBA and computed the flux distributions. For the comparisons, we computed two metrics: 1) the Pearson correlation between the predicted and measured intracellular fluxes, and 2) the average percentage error (see Materials and Methods) between the measured and predicted fluxes. A good prediction requires a noticeable correlation and a small average percentage error.

The results of the first set of experimental data [20] (pgm *vs.* Ref, rpe *vs.* Ref, zwf *vs.* Ref, wt5 *vs.* Ref, and wt7 *vs.* Ref) showed a considerably improved flux prediction for the REMI-Gex, REMI-TGex, REMI-TGexM, and REMI-GexM models as compared to the GX-FBA method as indicated by Pearson correlation and average percentage error (Figure 3a). The GX-FBA and REMI-Gex methods predicted a similar flux correlation for the experimental fluxes for the pgi *vs.* Ref and gapC *vs.* Ref cases (Figure 3a). For the second set of experimental data [21] (Nox *vs.* Ref and ATPase *vs.* Ref), REMI-TGex predicted better correlation than REMI-Gex and GX-FBA, and the average percentage error of GX-FBA was higher than that of REMI-TGex and REMI-Gex (Figure 2b). On an average across all nine comparisons (excluding references) we found that the REMI-Gex method has 32% higher Pearson correlation coefficient compared to the GX-FBA method, which indicates a remarkable improvement in the flux prediction. Since the REMI methods use an additional objective that is the minimization of the sum of fluxes (see Materials and Methods), we modified GX-FBA to imply the minimization of the sum of fluxes as an objective to perform an unbiased comparison. This modified GX-FBA prediction agreed less with the experimental results than the REMI predictions (Suppl. Figure 1), meaning that REMI outperforms GX-FBA in terms of predictions. REMI also has two advantages over GX-FBA and other relative expression methods in that, first, we do not need to estimate a reference flux distribution a priori, because two flux distributions for two different conditions are obtained in the same optimization framework in REMI (see Materials and Methods), second, REMI enumerates alternative solutions at the MCS, providing a higher confidence when investigating and analyzing results. Generating two separate flux distributions for the two compared conditions allows REMI to be more suitable to study the differential flux analysis between two conditions, and the extensive enumeration of alternative solutions provides robustness and flexibility in the biological interpretations of the provided data.

**Figure 3.**
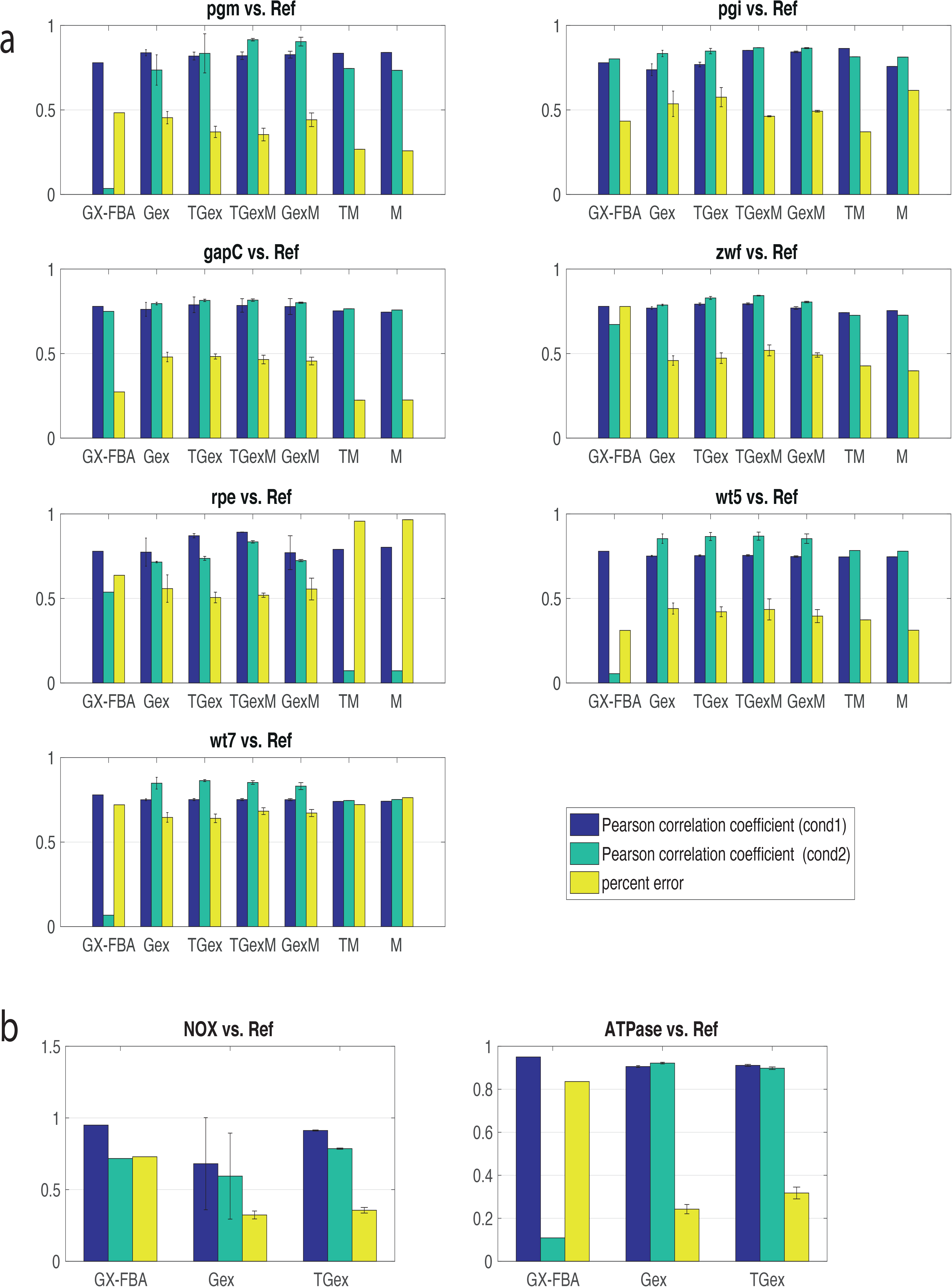
The comparison of steady-state fluxes between GX-FBA and REMI. The blue bar represents the Pearson correlation coefficient (PCC) between the experimental fluxes and predicted fluxes for the wildtype or reference state and the green bar represents the PCC for the mutant or overexpressed state. The third bar denotes the average percentage error between the experimental fluxes and predicted fluxes. Error bars (available only for the REMI method) represent the standard error of the mean of the alternative solutions. a) The comparison between the mutant and wildtype for the pgm, pgi, gapC, zwf, and rpe mutants as well as the comparison of the wildtype at growth rates of 0.5 hour^-1^ (wt5) and 0.7 hour^-1^ (wt7) compared to a reference wildtype with a 0.2 hour^-1^ growth rate (Ref). b) The comparison of NOX overexpression and ATPase overexpression against the wildtype.

Although all REMI methods were in relative agreement with the experimental fluxomic measurements, we did not observe a significant difference in the predicted results of REMI-Gex, REMI-TGex, and REMI-TGexM. However, as the fluxomic measurements were very limited around the central carbon metabolism, we cannot draw any overarching conclusions about the accuracy of REMI from these results, as this could only indicate that the major fluxomic differences occur in pathways outside of this one specific metabolic pathway. We believe that to investigate the differences in flux predictions across REMI methods, fluxomic and metabolomic measurements will be required on a grander scale, such as the genome level.

## Conclusions

We developed the computational tool, REMI, which combines gene-expression, metabolomics, and thermodynamics constraints with the mass balance constraints imposed in metabolic models to predict phenotypic changes in an organism upon environmental or genetic perturbations. As the integration of these three additional physiological constraint results in a highly reduced flexibility of the predicted feasible flux profiles, REMI enhances the quality of the computationally predicted fluxes. REMI’s novel formulation permits the extensive enumeration of alternative solutions because there exist several alternative sets of pathway that result in the same phenotype due to the complexity and interconnectivity of metabolic networks, meaning that the results provided by REMI more accurately reflect natural biological states than previously existing methods. The effectiveness of incorporating thermodynamic data with gene-expression and metabolomic data in reducing the flexibility of predicted feasible flux profiles. This means that we can obtain manageable set of physiological consistent hypothesis and physiological interpretations which have a higher confidence as they are consistent with a larger set of data. Applying REMI to experimental data has shown that there is not always a full consistency between gene-expression and metabolomic data, which shows that there is still much to learn about how gene expression and metabolism are linked. As systematic multi-omics integration remains a challenge, REMI opens the possibility of not only multi-omics integration, but also the identification of the crosstalk between the various omics present in a system.

## Materials and Methods

### Omics datasets and genome-scale model

Eleven total sets of experimental data that had been previously integrated into the genome-scale model (GEM) of *E.coli* by Kim *et al.* [23] and were originally obtained from two independent studies done by Ishii *et al.* (8 datasets) [20] and Holm *et al*. (3 datasets) [21] were used for the evaluation of the REMI methodology.

The three datasets from Holm *et al.* [20] comprise genome-wide transcriptomic data together with fluxomic data (21 measured fluxes) collected from three experimental conditions: wildtype *E. coli*, cells overexpressing NADH oxidase (NOX), and cells overexpressing the soluble F1-ATPase (ATPase). The eight datasets from Ishii *et al.* [20] include genome-wide transcriptomic, fluxomic (31 measured fluxes), and metabolomic (42 metabolites) data obtained under eight different experimental conditions: wildtype *E. coli* cells cultured at different growth rates of 0.2, 0.6, and 0.7 per hour along with single-gene knockout mutants of the glycolysis and pentose phosphate pathway (pgm, pgi, gapC, zwf, and rpe).

All analyses were performed using IJO1366, the latest GEM of *E. coli* [24]. The model comprises 2,583 reactions, 1,805 metabolites, and 1,367 genes. The REMI code is implemented in Matlab R2016a. Mixed-integer linear programming (MILP) problems were solved using the CPLEX solver on an Intel 12-core desktop computer running Mac.

### Integration of thermodynamic constraints into the genome-scale model

It has been previously shown that thermodynamic constraints not only effectively reduce the solution space of FBA by eliminating the thermodynamically infeasible fluxes from the solution space, but also allow the integration of metabolite concentrations. This provides important links between mass and energy balance and the phenotypic characteristics of the organism. The thermodynamic constraints, as depicted in Equation (1), were integrated into the IJO1366 model [3]. The standard Gibbs free energy 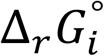 without corrections for the pH and ionic strength was estimated using the group contribution method [25].

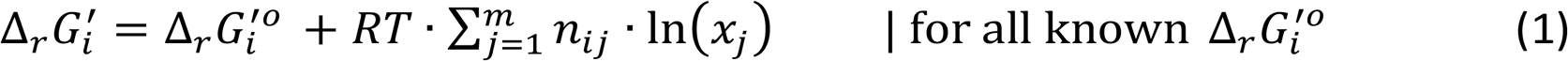

For each reaction of a GEM, the Gibbs free energy of the reaction 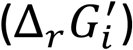 was computed, which considers the charge and the activity (*x*_*j*_) of each metabolite *j* given the pH, the metabolite concentration range, and the ionic strength at the cellular compartment where the reaction occurs.

### Assessment of tentative reaction flux ratios from the gene-expression and metabolomic data

We used the gene-protein-reaction (GPR) association rules acquired from the *E. coli* GEM to translate the relative gene-expression levels (being relatively up- or downregulated) to the differential and relative flux values of corresponding reactions. GPRs are not mapped as *one gene to one reaction,* meaning there are many cases in which one gene is mapped to several reactions and multiple genes are mapped to a single reaction, which are depicted with “and” and “or” affiliations, respectively.

In REMI, if the reaction R is associated with two genes (g1 “and” g2), the expression level ratios for genes g1 and g2 in the two corresponding conditions are calculated to obtain the geometric mean of the g1 and g2 ratios. Whereas, if the reaction R is associated with two genes (g1 “or” g2), the arithmetic mean of the obtained expression data ratios is calculated. Thus, from GPR associations, REMI computes the so-called tentative “reaction flux ratios” to further constrain the model. For the metabolomic data, the ratio of metabolite concentration for each metabolite (if available) is calculated for any two given conditions.

### Evaluating the differentially regulated metabolites and reactions

To evaluate whether a reaction or metabolite was up- or downregulated, the ratios (calculated as explained in the previous section) were sorted, with the top 5% (ratios > 1) selected as upregulated and the bottom 5% (ratios < 1) as downregulated using a conservative cutoff criterion, which is user-defined.

### REMI workflow

The REMI workflow along with an illustration of the method performed on a toy model is presented in Figure 4. REMI requires a GEM (FBA model) and sets of gene-expression and/or metabolomic data. The first step consists of data pre-processing wherein the FBA model is converted to a thermodynamic-based flux analysis (TFA) model [3] that incorporates the Gibbs free energy of metabolites and reactions into the model. The gene-expression/metabolite-level ratios are further systematically converted into reactions ratios to integrate them into the REMI methods. Based on the type of integrated data, there are three different REMI methods. REMI-TGex integrates thermodynamic and gene-expression data, REMI-TM integrates thermodynamic and metabolomic data, and REMI-TGexM integrates thermodynamic, gene-expression, and metabolomic data into an FBA model. Note that the REMI methods can be used without thermodynamic data, such as in REMI-Gex, which integrates gene-expression data into a FBA model. However, we will provide several examples to illustrate the power of thermodynamic constraints, when coupled with omics data, in reducing the predicted feasible flux profiles and better predicting the cell physiology.

**Figure 4:**
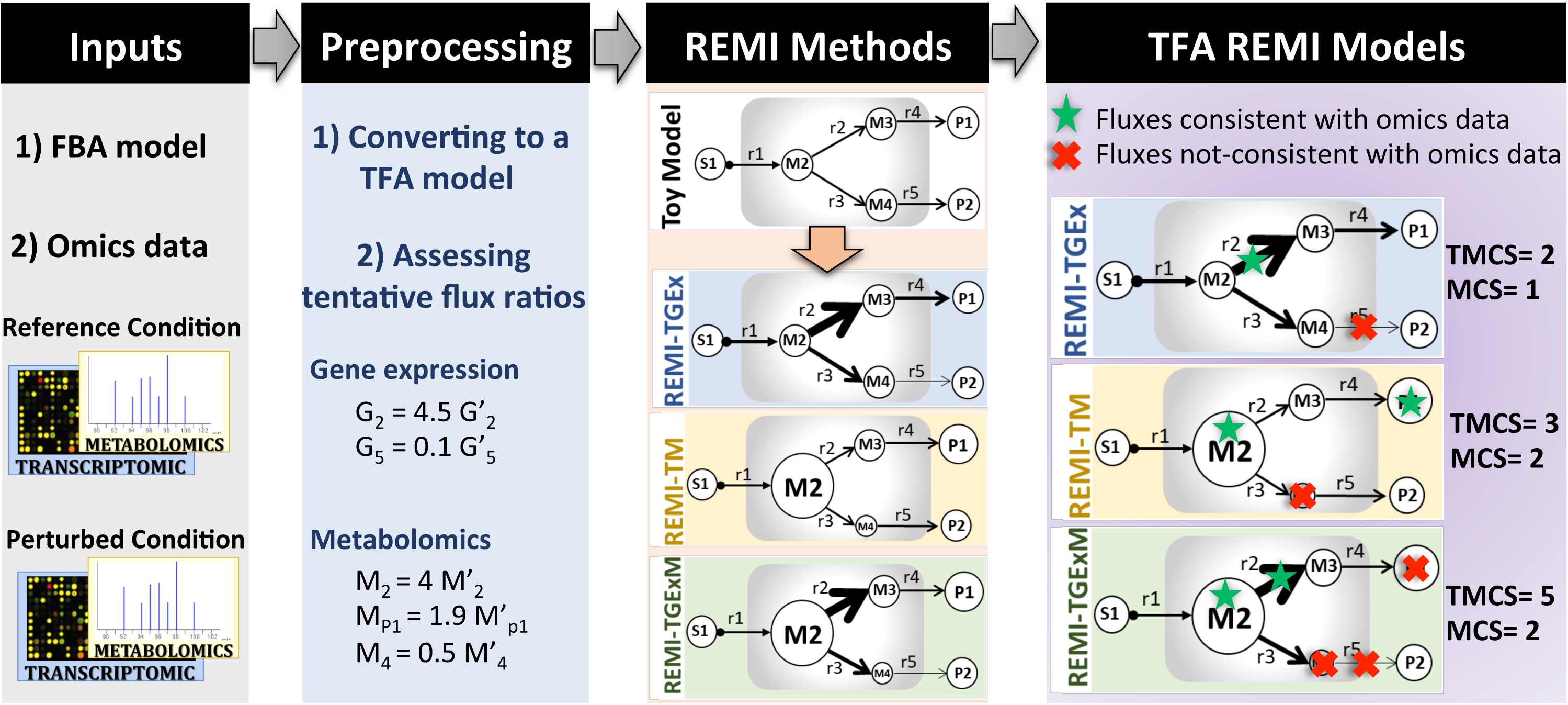
The REMI workflow requires two inputs: a genome-scale flux balance analysis (FBA) model and sets of gene-expression and/or metabolomic data. In the pre-processing step, the FBA model is converted into a thermodynamic-based flux analysis (TFA) formulation, and the relative flux ratios are further assessed based on the omics data. Also based on the omics data provided, REMI translates to the REMI-TGex, REMI-TM, and REMI-TGexM methods (third block). Examples of gene-expression and metabolomic data (second block) together with a toy mode (third block) are used to illustrate the applicability of the REMI methods. The theoretical maximum consistency score (TMCS) is the number of available omics data (for metabolites, genes (reactions), or both) and the maximum consistency score (MCS) is the number of those constraints that are consistent with fluxes and could be integrated into REMI models. The MCS is always equal to or smaller than the TMCS.

### Integration of relative gene-expression levels as constraints into a model (REMI-Gex)

For a given metabolic network that includes *R* reactions and *M* metabolites, bidirectional reactions are decomposed into forward and backward reactions to allow all fluxes to have positive values. Assuming that S is a stoichiometry matrix, S_mr_ is the stoichiometric coefficient associated with the metabolite m (m = 1,…, M) in reaction r (r = 1,…, R). Positive and negative stoichiometric coefficients of metabolites signify the substrate or products of a reaction. A binary variable z_r_ was assigned to each reaction r to ensure a positive flux v_r_ (Equation (2)) through the reaction r, and when z_r_ = 0, there was no flux. An additional constraint was formulated using Equation (3) to ensure that only one reaction directionality could be active and carry flux. α and β indicate the forward and reverse directions of a reaction.

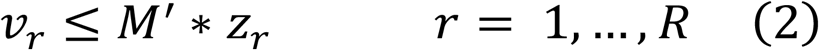

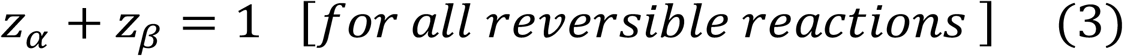

In REMI, two models are described for each given condition. Throughout the manuscript, the terms “wildtype” and “mutant” are used to better differentiate between the two conditions (or models) when describing the REMI framework. REMI can, however, be used for any two given conditions and is not restricted to the wildtype and mutant labels. Equation (4) specifies the mass balance constraints for the wildtype and mutant conditions at the steady state.

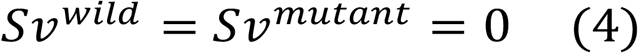

The relative information about the gene-expression levels or metabolite levels between the two given experimental conditions was formulated as additional constraints and integrated into the two representative models of the conditions. To do this, binary variables for the up- and downregulated reactions were assigned as *u* and *d,* respectively, where *n* is the total number of up- and downregulated reactions. For the upregulated reactions, a higher flux was enforced in the mutants as compared to the wildtype, while for downregulated reactions, a higher flux was enforced for the wildtype as compared to the mutant.

For *u* upregulated and *d* downregulated reactions, a total of *n* binary variables were generated (B_1_,…B_i_,…B_n_), where B*_i_* = 1 indicates the up- or downregulation of a reaction. Next, *n* constraints (Equations (6 and 7)) were added to enforce a basal flux in both the wildtype and mutant conditions. For *u* upregulated reactions, constraints (Equation (8)) were added to ensure a mutant flux could be higher 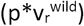 than a wildtype flux, where *p* is a reaction ratio between the wildtype and mutant (computed from gene-expression ratio). Constraints were added (Equation (9)) for *d* downregulated reactions that ensured a mutant flux was lower compared to a wildtype flux. In Equation (10), *n* constraints were added to form the boundary for the slack variables that are used in Equations (8) and (9), where *ε* = 10^-5^, *M*′ = 1000.

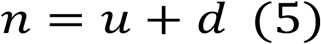

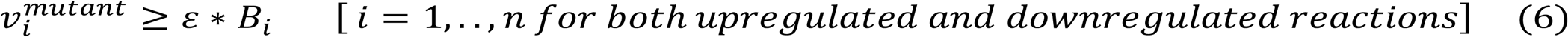

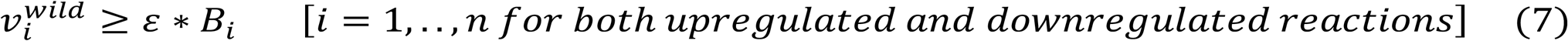

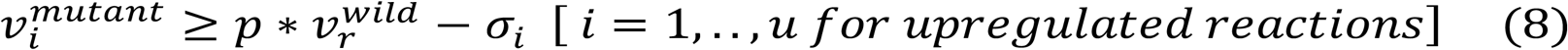

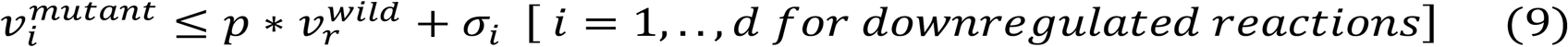

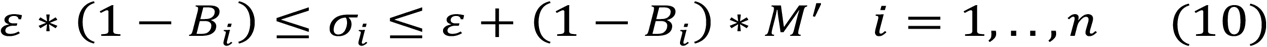

### Integration of relative metabolite levels as constraints into the model (REMI-M)

In GEMs, gene-level perturbations can mediate both reactions and their subsequent metabolites. Available studies show a correlation between gene changes and metabolite changes and infer that perturbations at the metabolite level are formed from perturbations in genes or reaction levels [26, 27]. Thus, if experimental evidence shows remarkable changes in a given metabolite abundance level across two conditions, the assumption is that there is an imbalance in the incoming or outgoing fluxes around that metabolite.

If the experimental data indicates that a metabolite is upregulated, it is assumed in REMI that either the sum of production 𝜙_*p*_ in condition 2 is greater than the 𝜙_*p*_ in condition 1 or the sum of consumption 𝜙_*c*_ in condition 2 is less than the 𝜙_*c*_ in condition 1 (Figure 5b). Due to mass balance, 𝜙_*p*_ and 𝜙_*c*_ will be equal.

**Figure 5.**
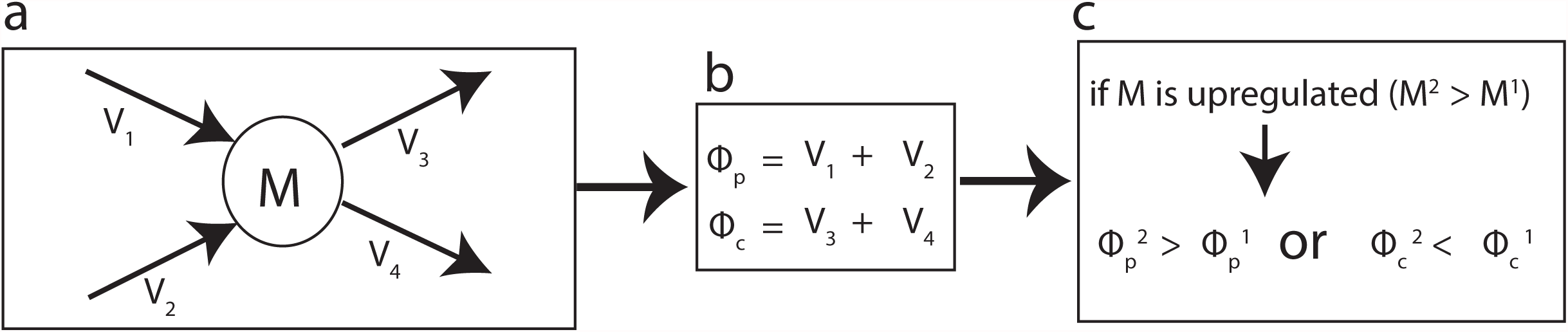
The relative metabolite integration within REMI.

In Equation (11), the sum of production and of consumption of a metabolite *i* is shown, where the metabolite is produced by reactions 1 and 2 and is consumed by reactions 3 and 4 (Figure 5a).

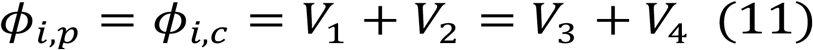

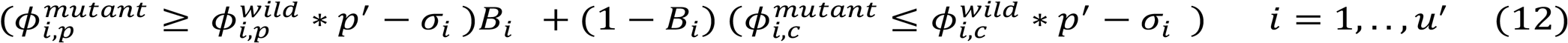

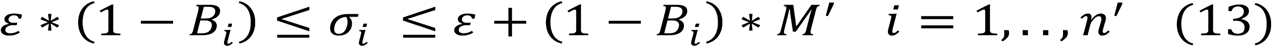

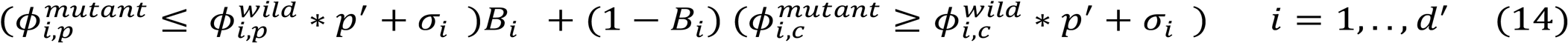

Based on available experimental measurements of metabolite abundance, REMI finds the total number (*n’*) of up- and downregulated metabolites, where *u’* and *d’* are up- and downregulated metabolites, respectively. For an up-regulated metabolite *i* (i.e. in the mutant vs. wildtype), either more production or less consumption is enforced in the mutant compared to the wildtype using Equations (12) and (13). In Equation (12), a binary variable (*B*_*i*_) is introduced, which switches to production if *B*_*i*_ = 1 and to consumption if *B*_*i*_ = 0. Similarly, for downregulated metabolites *i*, less production or more consumption is enforced in the mutant compared to the wildtype (Equations (13) and (14), see supplementary description for more detail).

### The objective function and the consistency score based on the relative expression data

Based on the assumption that alterations in gene-expression or metabolite levels within two different physiological conditions results in differential flux profiles, REMI defines such alterations as constraints and integrates them accordingly into the two metabolic models corresponding to the two conditions. However, as additional constraints reduce the solution space of FBA, particularly in the case of multi-omic data integration, the resulting models might not be feasible. Therefore, the objective function (Equation (15)) was formulated in such a way as to obtain feasible models with a maximum agreement between the relative expression and metabolite levels and their corresponding constraints. Equation (15) maximizes the agreement with experimental data using mathematical optimization principles subject to Equations (5)–(10), where n is the total number of up- and downregulated reactions. The maximum consistency score (MCS) is the sum of the binary variables (Equation (15)) in the outcome of the optimization that is formulated in REMI.

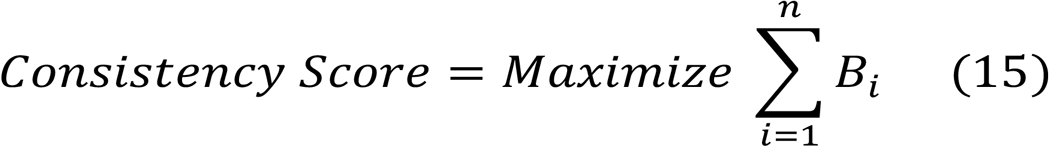

### Alternative enumeration for consistency score

An aforementioned mathematical optimization model (objective function (15) subject to Equations 1–9) allows us to maximize the total number of consistent reactions between the differential gene-expression or metabolite levels with the differential flux profiles between two models and to obtain a maximum consistency score (MCS). Depending on the flexibility of the model, many alternative flux distribution profiles for a given MCS, and subsequently MCS-n, are possible. MCS and MCS-n represent optimal and suboptimal consistency, respectively. To enumerate alternative solutions, integer cut constraints (Equation (16) [28] were used as follows:

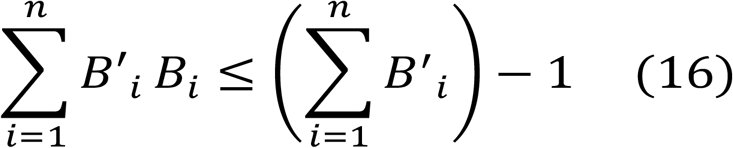

The left-hand side of Equation (16) determines the number of up- and downregulated reactions in the current solution that carries fluxes in the first MCS solution. The right-hand side represents the number of reactions that carry fluxes in MCS-1. The inequality ensures that the new solution differs at least by one new reaction that carries flux compared to the previous solution. Repeating this procedure allows the enumeration of alternative solutions for each MCS.

### Combined consistency score using relative expression and relative metabolite (REMI-GexM)

To concurrently integrate both the relative gene-expression data and the relative metabolite levels, an integrated mathematical optimization model was built with a global objective function (Equation (17) subject to a combined set of constraints, i.e. equations (5)–(14). This optimization model was then solved to maximize the objective, which is the combined consistency score of the two sets of constraints.

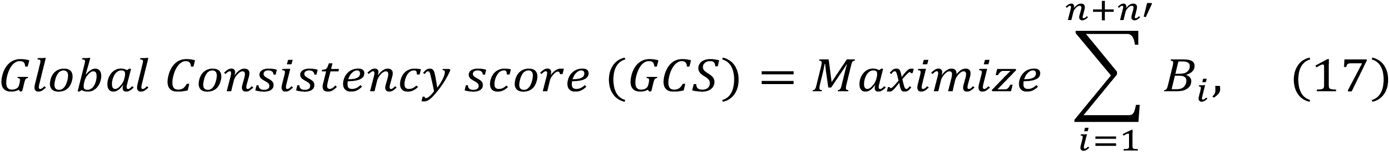

where *n* and *n’* represent a total number of up- and downregulated metabolites and up- and downregulated genes, respectively.

### Representative flux distribution profiles

To compare the REMI-predicted fluxes with the experimentally measured ones, predicted flux distribution profiles were required. To obtain such predicted flux profiles, all the alternative solutions at MCS were first enumerated. Then, an additional optimization was performed by minimizing the sum of the fluxes for each alternative solution to obtain a representative flux profile for benchmarking REMI against the experimental flux measurements.

### Metrics for comparing the predicted *in silico* fluxes with experimentally measured fluxes

To effectively compare the predicted *in silico* fluxes from REMI with the corresponding ^13^C-determined *in vivo* intracellular fluxes, the following two metrics were used: the uncentered Pearson correlation coefficient (Equation 18), and the average percentage error in predicted fluxes (Equations (18)-(20)). The uncentered Pearson correlation is a good metric for the flux comparison, as fluxes are usually not centered, and it has been used for comparing two flux vectors [23].

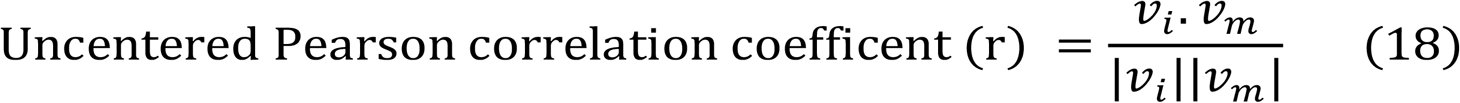

In Equation (18), *v*_*i*_ and *v*_*m*_ are the *in silico* and measured vectors of the fluxes, respectively. The correlation coefficients +1 and -1 indicate a strong positive and negative linear relationship between *v*_*i*_ and *v*_*m*_, and the 0 correlation coefficient indicates no linear relationship between v_i_ and v_m_.

The average percentage error has been used in the GX-FBA method [12] to compare two fluxes. In Equation (19), the d_r_ is used to measure the relative deviation between the two fluxes in two conditions, where *x* and *y* correspond to the flux of a given reaction in condition 1 and condition 2, respectively. Since |*d*_*r*_| lies between 0 and 1, one can consider d_r_ as a percentage flux change from condition 1 to condition 2. The average (per reaction) percentage error, *e*, in the predicted *in silico* fluxes was calculated using Equation (20), where 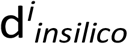 and 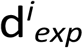 indicate relative deviation in predicted *in silico* flux using methods such as REMI and GX-FBA, and experimentally measured flux and N represent the number of reactions with available experimental flux data.

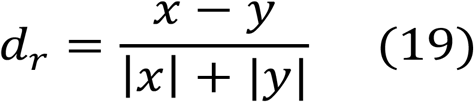

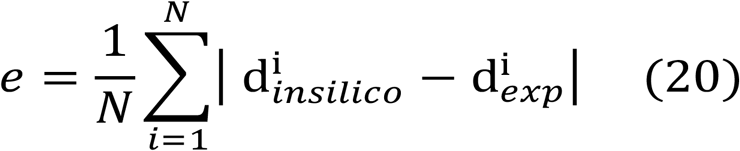

### Assessing the relative flexibility of metabolic systems

For a given system, the FBA results in a solution space of optimal flux profiles, and the magnitude of this solution space indicates the metabolic flexibility of the system. The integration of the thermodynamic knowledge of reactions as well as condition-specific experimental data, e.g. gene-expression or metabolomic data, constrains the metabolic system to a less flexible one. Thus, the solution space and the subsequent range of the metabolic responses are reduced. Comparing and quantifying the relative flexibility of a metabolic system before and after constraint is a decent indication of the effectiveness of the data integration [29]. Performing a flux variability analysis (FVA) outlines the flux variability range of each reaction in the system for the two conditions as follow:

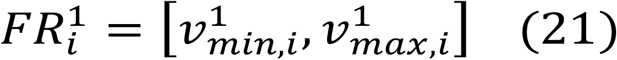

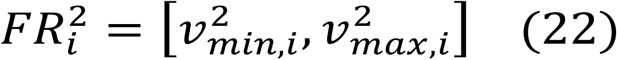

The *relative flexibility* (*RF*) for reaction *i* is calculated using the following equation:

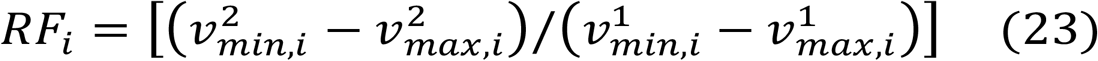

where 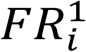 and 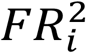 represent the flux variability range of reaction *i* at each of the two conditions, one condition is usually designated as a reference condition or reference state, such as when comparing the relative flexibility of a metabolic system with (condition 1) and without (condition2) thermodynamic constraints. The value of RF that is computed for each reaction *i* reflects the relative changes in the flux variability range of one condition compared to the other condition. The *global relative flexibility change between two given condition* is then computed by averaging the F_i_ values for each reaction *i* that carry flux in the reference state.

## Supporting information

## Supplementary Tables

Table S1. Flux variability ranges for 45 reactions at different consistency score.

Table S2. Relative flexibility of metabolic subsystems for 9 different comparisons.

## Supplementary Figures

Figure S1. Relative flexibility over all reactions for 9 different comparisons.

Figure S2. Comparison of the flux perdition between the GX-FBA and REMI-Gex method.

## Acknowledgments

VP and VH are supported by the RTD grant MalarX within SystemsX.ch, the Swiss Initiative for Systems Biology evaluated by the Swiss National Science Foundation. VP and VH are supported by the École polytechnique fédérale de Lausanne. NH is supported through the RTD grant MicroScapesX, no. 2013/158, within SystemX, the Swiss Initiative for System Biology evaluated by the Swiss National Science Foundation.

## Author Contributions

Conceptualization and methodology: VP, NH, and VH. Implementation of the method and data analysis: VP. Wrote the paper: VP, NH, and VH. Supervision: VH.

