## Supplementary material for "Enhanced flux prediction by integrating relative expression and relative metabolite abundance into thermodynamically consistent metabolic models"

Supplementary Information

**Integration of relative metabolite changes**

For integrating metabolites with the REMI method, we used the following equations that are also described in main text. Based on available experimental measurements of metabolite abundance, REMI finds the number of up- (*u’*) and downregulated (*d’*) metabolites. For an up-regulated metabolite *i* (i.e. in the mutant vs. wildtype)*,* either more production or less consumption is enforced in the mutant compared to the wildtype using Equations (1) and (2). In Equation (1), a binary variable (*B_i_*) is introduced, which switches to production if *B_i_* = 1 and to consumption if *B_i_* = 0. Similarly, for downregulated metabolites *i*, less production or more consumption is enforced in the mutant compared to the wildtype (Equation (2)).

$${(\phi}_{i,p}^{mutant}\geq\phi_{i,p}^{wild}*p^{'}-\sigma_{i} )B_{i} +(1-B_{i}) {(\phi}_{i,c}^{mutant}\leq\phi_{i,c}^{wild}*p^{'}-\sigma_{i} ) i=1,..,u^{'} (1)$$

$${(\phi}_{i,p}^{mutant}\leq\phi_{i,p}^{wild}*p^{'}+\sigma_{i} )B_{i} +(1-B_{i}) {(\phi}_{i,c}^{mutant}\geq\phi_{i,c}^{wild}*p^{'}+\sigma_{i} ) i=1,..,d^{'} (2)$$

Since equations 1 and 2 comprise terms that are a multiplication of binary and continuous variables. For example, in equation (1) multiplication terms are: ${(\phi}_{i,p}^{mutant}*B_{i}$ ), ${(\phi}_{i,p}^{mutant}*B_{i})$, ${(\phi}_{i,c}^{mutant}*B_{i}), {(\phi}_{i,c}^{wild}*B_{i})$, and $(\sigma_{i} *B_{i})$. For each multiplication term we add three additional constraints to change nonlinear terms into linear terms. Assume we take an expression *Z*= *C***B*, where C is a continuous variable which value is a positive value and B is a binary variable. For each expression (*Z*= *C***B*) we added constraints (equations (3)-(5)). M is a large integer number.

$$Z\leq M*B (3)$$

$$Z\leq C (4)$$

$$Z\geq C-\left( 1-B \right)M (5)$$

If *B* is zero then equation (3) and (5 ) ensures that Z will be negative number or zero and if B is 1 then equations (3)-(5) ensures Z is equal to C.
