## Supplementary figures and images for "Enhanced flux prediction by integrating relative expression and relative metabolite abundance into thermodynamically consistent metabolic models"

### Supplementary file 4

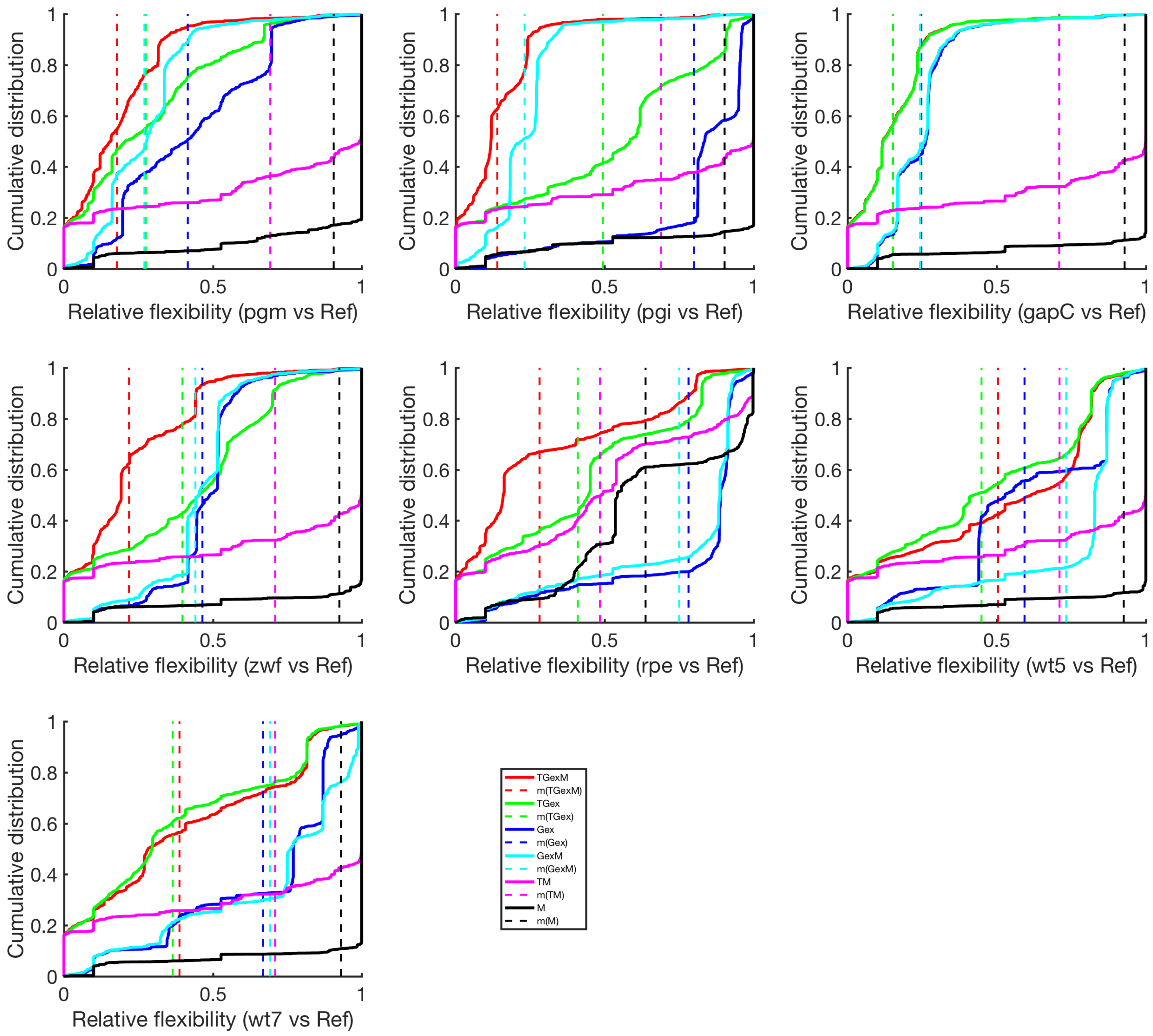

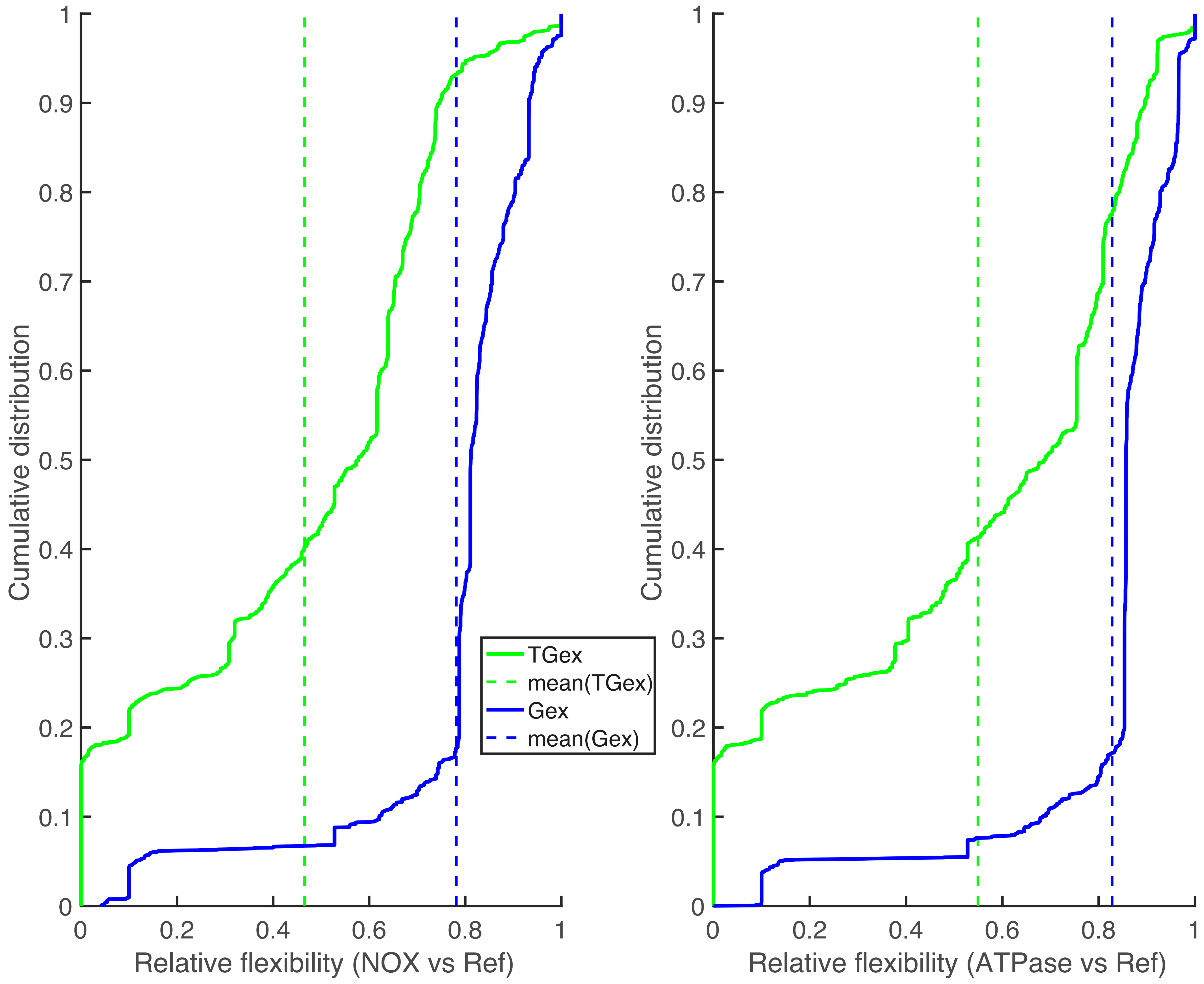

### Supplementary file 5

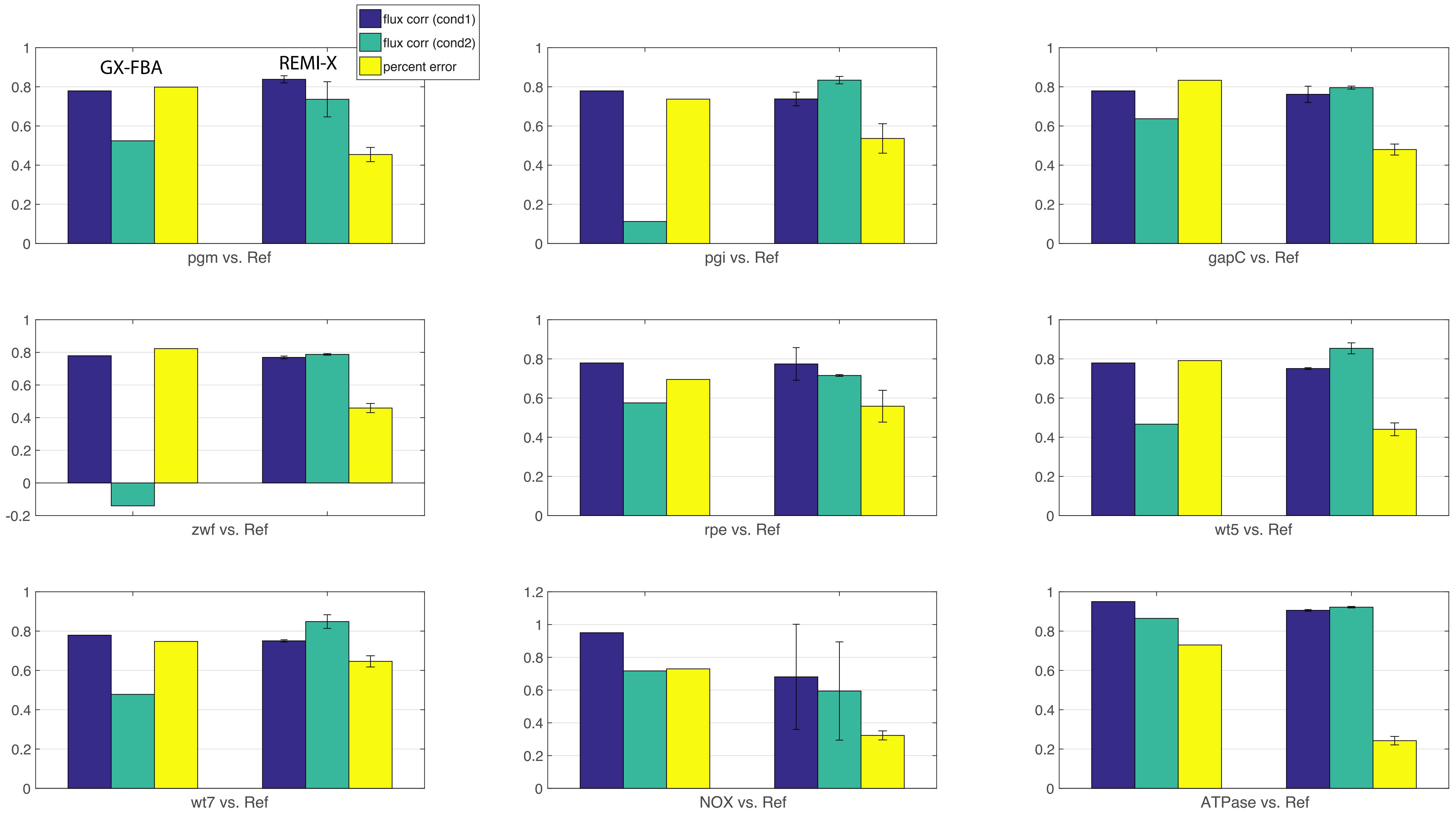
